# DNA methylation and demethylation underlie the sex difference in estrogen receptor alpha in the arcuate nucleus

**DOI:** 10.1101/2021.03.25.437030

**Authors:** Laura R. Cortes, Carla D. Cisternas, Iagn N.K.V. Cabahug, Damian Mason, Emma K. Ramlall, Alexandra Castillo-Ruiz, Nancy G. Forger

## Abstract

**Introduction:** Neurons expressing estrogen receptor (ER) α in the arcuate nucleus (ARC) of the hypothalamus sex-specifically control energy homeostasis and bone density. Females have more of these neurons than do males, but how this sex difference develops is unknown.

**Objective:** We tested the hypothesis that DNA methylation and/or demethylation control the development of a sex difference in ERα in the ARC.

**Methods:** ERα immunoreactive neurons were quantified at birth and at weaning in male, female and testosterone-treated female mice that received neonatal, intracerebroventricular injections of vehicle or zebularine, a DNA methyltransferase inhibitor. Methylation status of *Esr1* was determined in the ARC and ventromedial hypothalamus (VMH) using bisulfite conversion of DNA followed by pyrosequencing. Small interfering RNAs against ten-eleven translocases were used to examine effects of demethylation on ERα cell number.

**Results:** A sex difference in ERα cell number in the ARC, favoring females, developed between birth and weaning and was due to programming effects of testosterone. Zebularine treatment eliminated the sex difference in ERα in the ARC at weaning by *decreasing* ERα *in females* to male-like levels. Previously, the same treatment *increased* ERα *in males* in the VMH. A promoter region of *Esr1* exhibited sex differences in opposite directions in percent of total methylation in the ARC (females > males) and VMH (males > females). Moreover, neonatal inhibition of demethylation increased ERα in the ARC of males.

**Conclusion:** DNA methylation and demethylation regulate ERα cell number in the ARC, and methyl marks may paradoxically activate *Esr1* in this region.

## Introduction

A transient exposure to testosterone during perinatal life underlies the emergence of many sex differences in the mammalian brain [1,2]. For example, male mice have more neurons expressing calbindin and vasopressin in several forebrain regions [3,4], while females have more neurons expressing estrogen receptor (ER) α [5,6]. In some cases, sex differences in developmental cell death is prevented, *e*.*g*., [7,8]. In the latter case, sexual differentiation is often a process of establishing stable, sex-specific patterns of gene expression in neurons, *i*.*e*., the differentiation of neurochemical phenotype. Although sex differences in neurochemical phenotype are common, the underlying molecular processes are largely unknown.

The arcuate nucleus (ARC) is located in the mediobasal hypothalamus and exhibits more ERα expressing neurons in females than in males, at least in rats [6]. ERα neurons in the ARC play important roles in the control of feeding behavior, energy homeostasis, and the regulation of bone density [9–11], but what causes females to have more of these neurons is unknown.

Epigenetic modifications, such as DNA methylation, play prominent roles in the differentiation of cell type throughout the body and may underlie sexual differentiation of ERα neurons [12–16]. For example, we recently found that inhibiting DNA methylation in newborn mice reduces or eliminates the usual female bias in the number of neurons expressing ERα in the preoptic area and ventrolateral area of the ventromedial nucleus of the hypothalamus (VMHvl), which lies adjacent to the ARC [14,16].

DNA methylation is catalyzed by DNA methyltransferases (DNMTs) and usually occurs at the 5th carbon position of cytosine residues followed by a guanine (CpG sites) [17]. DNA methyl marks (5-methylcytosine; 5mC) can be quite long-lasting [18], but they can also be removed through a series of oxidative steps catalyzed by ten-eleven translocation enzymes (TETs). In the first step, 5mC is converted to 5-hydroxymethylcytosine (5hmC), which is both an intermediary in demethylation and an independent epigenetic mark in its own right [19]. Although a growing number of exceptions are reported [20–25], 5mC is typically associated with gene repression, whereas 5hmC is most often associated with gene expression [22,26–28]. Thus, the emergence of a particular cell phenotype in part depends on the balance between methylation (controlled by DNMTs) and demethylation (controlled by TETs) at specific genes.

The methylation landscape of the brain changes dynamically during the first few weeks (rodents) or years (humans) of life [29]. We recently found that the expression and enzyme activity of DNMTs and TETs peak in the mouse hypothalamus during the first postnatal week [30], which coincides with the critical period for sexual differentiation of the brain and behavior. Furthermore, we found sex differences in *Tet* gene expression in the neonatal hypothalamus, with greater expression of *Tet2* and *Tet3* in males [30]. Here, we manipulated DNMTs and TETs neonatally to test the hypothesis that DNA methylation and/or demethylation underlie sexual differentiation of ERα in the ARC. We also examined DNA methylation of *Esr1* promoter regions in both the ARC and neighboring VMHvl.

## Materials and Methods

### Animals

C57BL6/J mice were bred in our vivarium and checked daily for pups. Mice were housed in cages with corn cob bedding in a room maintained at 22°C with food and water available *ad libitum*. Animal procedures were performed in accordance with the National Institutes of Health animal welfare guidelines and were approved by the Georgia State University Institutional Animal Care and Use Committee.

### Neonatal testosterone treatment and DNA methyltransferase inhibition

The brains used in this portion of the study were the same as those used in a prior study reporting on the effects of a neonatal inhibition of DNA methylation on ERα in the POA and VMHvl [16]. Male and female controls received a subcutaneous injection of 25 µL of peanut oil on the day of birth and following day, while androgenized females received testosterone propionate (100 µg in 25 µL peanut oil; Millipore Sigma, St Louis, MO). Half of the animals in each of the three hormonal groups also received intracerebroventricular (ICV) injections of either vehicle or a DNA methyltransferase inhibitor, zebularine, on the day of birth (postnatal day (P) 0) and P1. Vehicle (500 nL per hemisphere; 10% dimethyl sulfoxide in 90% saline) and zebularine (300 ng in 500 nL vehicle; Millipore Sigma) were injected into the lateral ventricles of cryoanesthetized pups, as described previously [16]. All animals were tattooed for identification and were sacrificed prior to puberty at P25. This age was chosen because it allowed us to examine relatively long-lasting effects of the neonatal inhibition of DNA methylation in the absence of confounding effects of post-pubertal gonadal hormones.

### Neonatal *Tet* inhibition

*Tet* mRNA expression in the hypothalamus is elevated during the first postnatal week, and sex differences in *Tet2* and *Tet3* expression (male > female) are observed during this time [30]. To determine the role of demethylation in the development of the sex difference in ERα expression in the ARC and VMHvl, we used small interfering RNAs (siRNAs) to reduce *Tet* expression in males to female-like levels. ICV injections were performed on cryoanesthetized pups as described above. On P5, females received a control injection of non-targeting RNAs in delivery media while males received either the control or a cocktail of SMARTPool siRNAs targeted against *Tet2* and *Tet3* (400-500 pmol; Accell, Horizon Discovery, Cambridge, UK). Each SMARTPool contains a mixture of four siRNAs targeting the same mRNA. Previous studies using Accell siRNAs report a neuron-selective knock-down that is maximal 2-4 days after a single ICV injection [31,32]. Thus, a subset of brains was collected two days after injection (P7), and punches were taken of the anterior and posterior hypothalamus to confirm gene knock-down by RT-qPCR, as described in [30]. The remaining brains were harvested at P25, immunohistochemically stained for ERα, and the ARC and VMHvl were analyzed as described below.

### Immunohistochemistry and quantification of ERα

Brains were collected and fixed in 5% acrolein in 0.1 M phosphate buffer for 24 h, then transferred to 30% sucrose in 0.1 M phosphate buffer for several days. Brains were frozen-sectioned into four series (zebularine-treated animals) or two series (siRNA-treated animals) of 30 μm thickness and stored in cryoprotectant (30% sucrose, 30% ethylene glycol, and 1% polyvinylpyrrolidone in 0.1 M phosphate buffer) until immunohistochemical staining. One series was stained for ERα (rabbit anti-ERα, 1:20,000; EMD Millipore, Billerica, MA) as described previously [16]. A separate cohort of P1 untreated tissue was also immunostained for ERα to determine if sex differences were present at this age.

Images through the ARC were captured using Stereo Investigator software (MBF Bioscience, Williston, VT) and ERα labeling was analyzed using ImageJ (Version 1.52/1.53; National Institutes of Health, Bethesda, MD). The boundaries of the ARC were traced and the number of pixels above threshold was quantified per μm^2^. Pixels above background were automatically determined using the Shanbhag algorithm, and the four sections through the ARC with the greatest labeling were summed for each animal. To validate this method, we also manually counted ERα cells in a subset of male and female animals using Stereo Investigator and confirmed a very similar pattern of results.

### Quantification of cell death

To determine whether zebularine treatment altered developmental cell death, brains from a separate cohort of vehicle- and zebularine-treated pups were collected 6 hours after the last injection on P1 and immunolabeled for activated caspase-3 (AC3) as previously described [33]. We drew contours around the ARC and counted the number of AC3-positive cells bilaterally. Cell death density was obtained by dividing the number of AC3-positive cells by the area sampled and is expressed as AC3 cells per mm^2^.

### Stereological Cell Counts

A separate series of brain sections from vehicle- and zebularine-treated females was stained with thionin to determine whether zebularine altered the total number of cells in the ARC at weaning. Unbiased stereology using the optical fractionator probe was performed using Stereo Investigator software (MBF Biosciences). Contours were drawn around the region of interest at low power, and counts of cells exhibiting a neuronal morphology were performed using the 100x oil objective within a 324 μm^2^ counting frame and using a 4,900 μm^2^ sampling grid. The Gundersen coefficient of error for counts was between 5-7% [34].

### Bisulfite mapping of *Esr1* in the ARC and VMHvl

A separate cohort of male (n = 84) and female (n = 84) pups received ICV injections of vehicle or zebularine on P0 and P1. Forty-eight vehicle-treated pups (24 of each sex) were sacrificed ∼4 h after ICV injection on P1, and all remaining animals were collected on P25. Brains were removed, flash-frozen in 2-methylbutane cooled to -20° C, and cut on a cryostat to the level of the ARC and VMHvl. Punches were taken from the ARC and VMHvl and kept at -80° C until processing. Because a punch of the ARC damages the adjacent VMHvl, and vice versa, separate animals were used for each brain region. Punches from three animals of the same sex were pooled per sample. Bisulfite conversion followed by pyrosequencing was performed by EpigenDx (Hopkinton, MA) to obtain single-nucleotide resolution of modified cytosines in *Esr1*. Specifically, we assessed the modification of cytosines in 16 CpG sites in the untranslated regions of Exons A and C. Because bisulfite sequencing cannot distinguish between 5mC and 5hmC, we refer to the results of this analysis as “total methylation” (as in [22,35]). Methylation of these regions was previously shown to associate with changes in *Esr1* gene expression in the mouse brain during development or after bisphenol A exposure [36,37].

### Statistics

Independent, two-tailed t-tests were used to compare ERα labeling in the ARC of untreated males and females on P1. Two-way ANOVA (group-by-treatment) was used to compare ERα labeling at P25 in males, females, and testosterone-treated females that received either vehicle or zebularine neonatally. Results of pyrosequencing were analyzed using a two-way repeated measures ANOVA for P1 samples, with sex as the between-subjects factor and CpG site as the repeated measure. Pyrosequencing at P25 was analyzed with a three-way repeated measure ANOVA with sex and zebularine treatment as between-subjects factors and CpG site as the repeated measure. A significant main effect of sex was followed by post-hoc tests of individual CpG sites. Developmental changes in mean methylation across CpG sites were analyzed with two-way ANOVAs (age-by-sex). Effect of *Tet* siRNA treatment on ERα expression was analyzed by one-way ANOVA. Fisher’s LSD’s post-hoc or Tukey’s tests were used where appropriate and all analyses were performed using Prism Version 9 (GraphPad Software, San Diego, CA).

## Results

### A sex difference in ERα cell number in the ARC is programmed by neonatal testosterone and eliminated by an inhibition of DNA methylation

We observed prominent ERα labeling in the ARC of untreated animals that appeared confined to cell nuclei (Figure 1). There was no sex difference at P1 (Fig. 1a,c), but females had more ERα labeling than males at weaning (P25; t_16_ = 3.64, *P* = 0.002; Fig. 1b,d). Thus, the sex difference in ERα in the ARC previously reported in adult rats [6] is present prior to puberty in mice. To examine hormonal control of the sex difference, we compared ERα in the ARC at P25 in the males, females, and females treated neonatally with testosterone from our previous study [16]. As expected, control females had greater ERα labeling than did control males (*P* < 0.0001 for Vehicle Female vs Vehicle Male; Fig. 2). Neonatal treatment of females with testosterone markedly reduced ERα labeling at P25 (*P* < 0.0001 for Vehicle Female vs Vehicle Female + T), indicating that the sex difference is due to programming effects of testosterone. Our testosterone treatment, in fact, hyper-masculinized females relative to vehicle-treated males (*P* < 0.01).

**Fig. 1.**
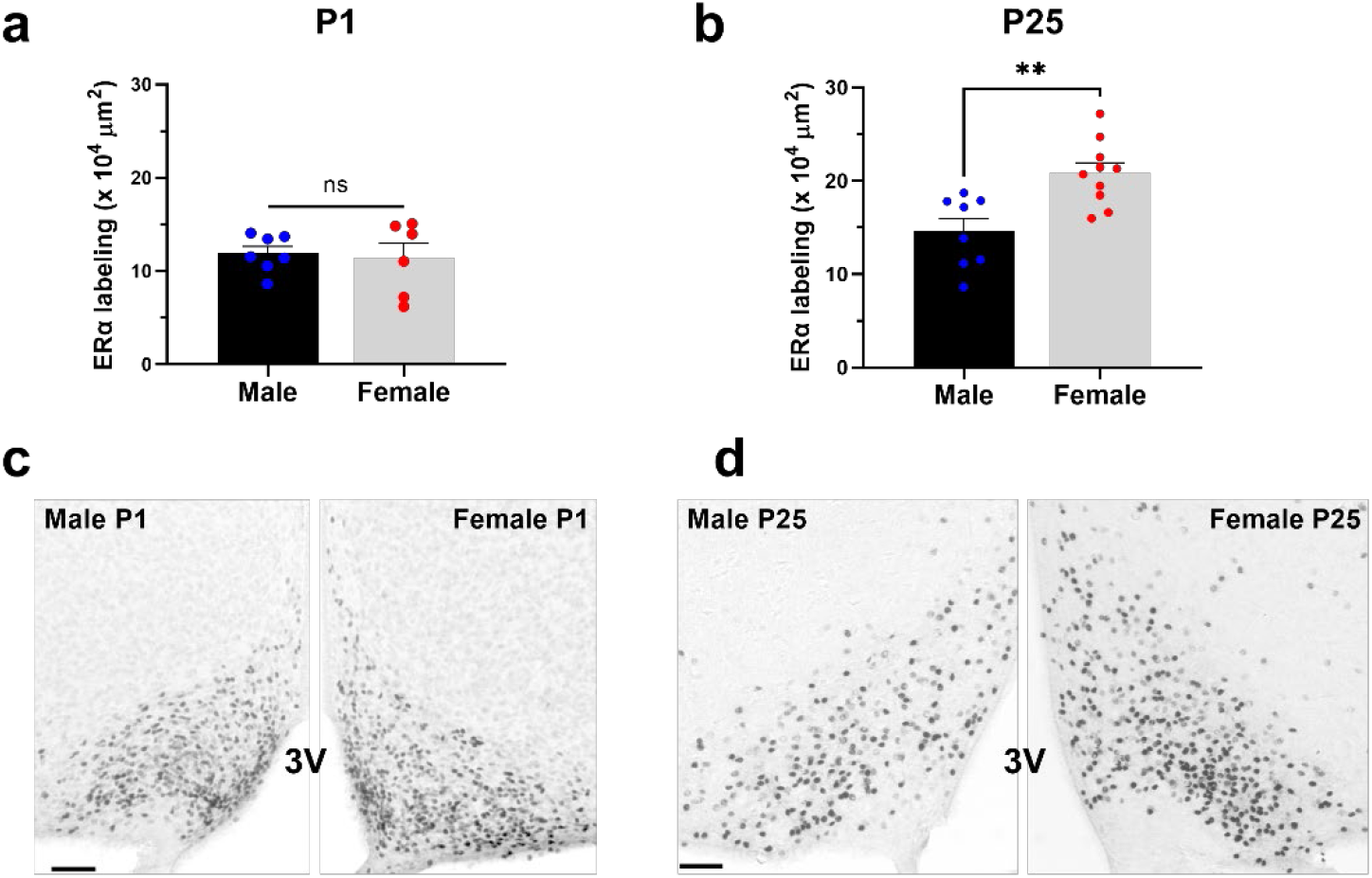
A sex difference in ERα labeling in the ARC emerges at weaning. (a,c) Males and females have similar ERα labeling one day after birth (P1). (b,d) Females have more ERα labeling than males at P25. Mean + standard error of the mean and individual data points are depicted. Scale bar = 50 µm. 3V = third ventricle.

**Fig. 2.**
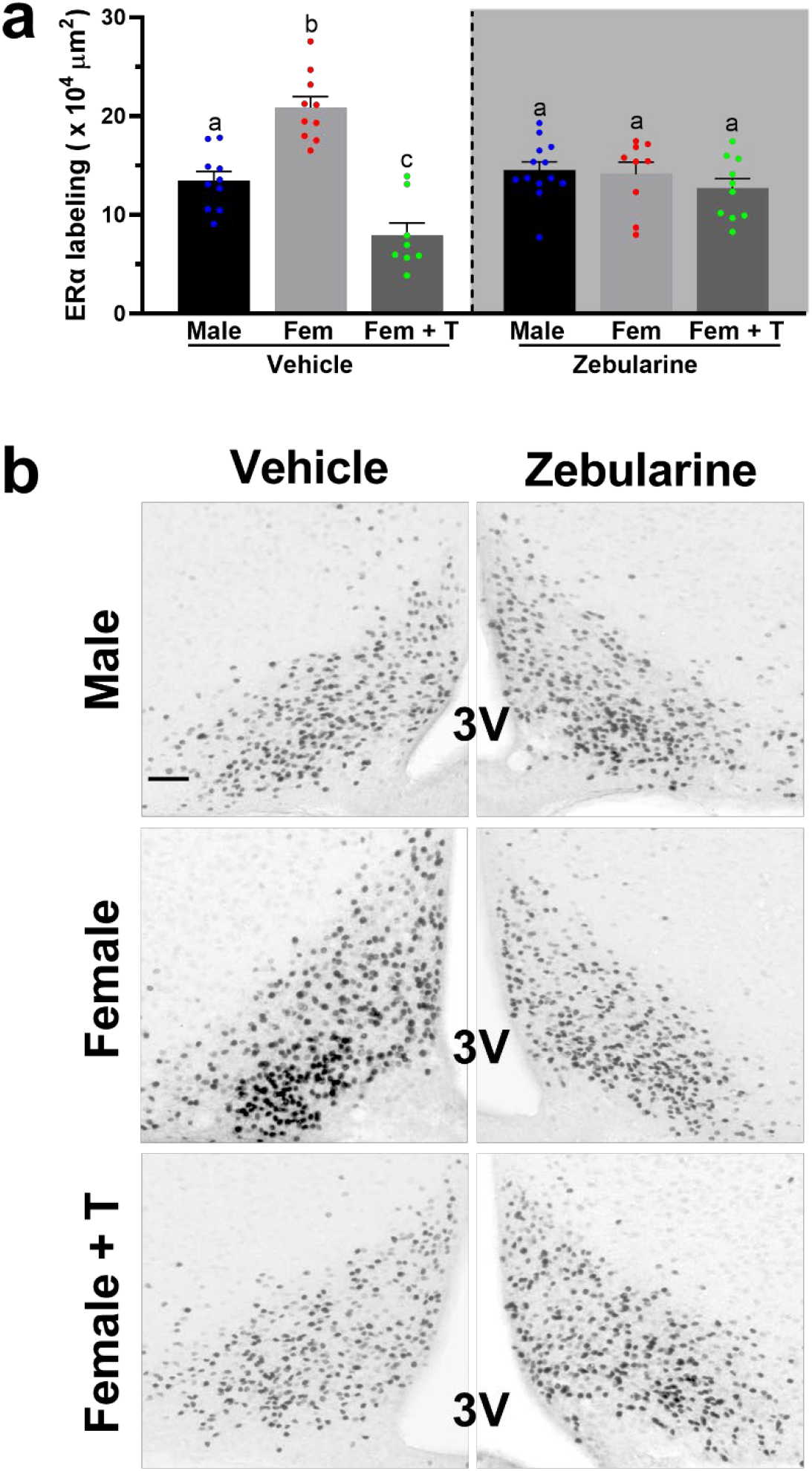
The sex difference in ERα labeling in the ARC is programmed by neonatal testosterone and is eliminated by a neonatal inhibition of DNA methyltransferases. (A) Mean ERα labeling at P25 in control males and females and in females treated with testosterone (T) at birth. Animals also received ICV injections of vehicle (left) or zebularine (right) during the first two days of life. Bars marked by different letters are significantly different (*P* < 0.01). (B) Photomicrographs showing ERα labeling in the six groups. Scale bar = 50µm.

To probe the role of DNA methylation, we compared ERα labeling in the ARC in animals treated neonatally with zebularine or vehicle. There was no main effect of neonatal zebularine treatment, but a highly significant interaction between group (Male, Female, and Female +T) and treatment (Vehicle vs. Zebularine; F_2,54_ = 15.12, *P* < 0.0001; Fig. 2). Neonatal zebularine treatment decreased ERα labeling in control females (*P* < 0.0005) to male-like levels, while having no effect on males. Zebularine also slightly increasing labeling in testosterone-treated females (*P* = 0.035). As a result, group differences in ERα labeling in the ARC were eliminated among the zebularine-treated animals (Fig. 2). The same pattern of results was obtained using direct cell counts of ERα, rather than automated analyses of labeling (not shown). The effects of zebularine were independent of a change in developmental cell death in both sexes at P1 (Fig. 3a), or total cell number in females at P25 (Fig. 3b). Thus, neonatal inhibition of DNA methylation reduced the proportion of neurons that express ERα in females, rather than the number of surviving neurons.

**Fig. 3.**
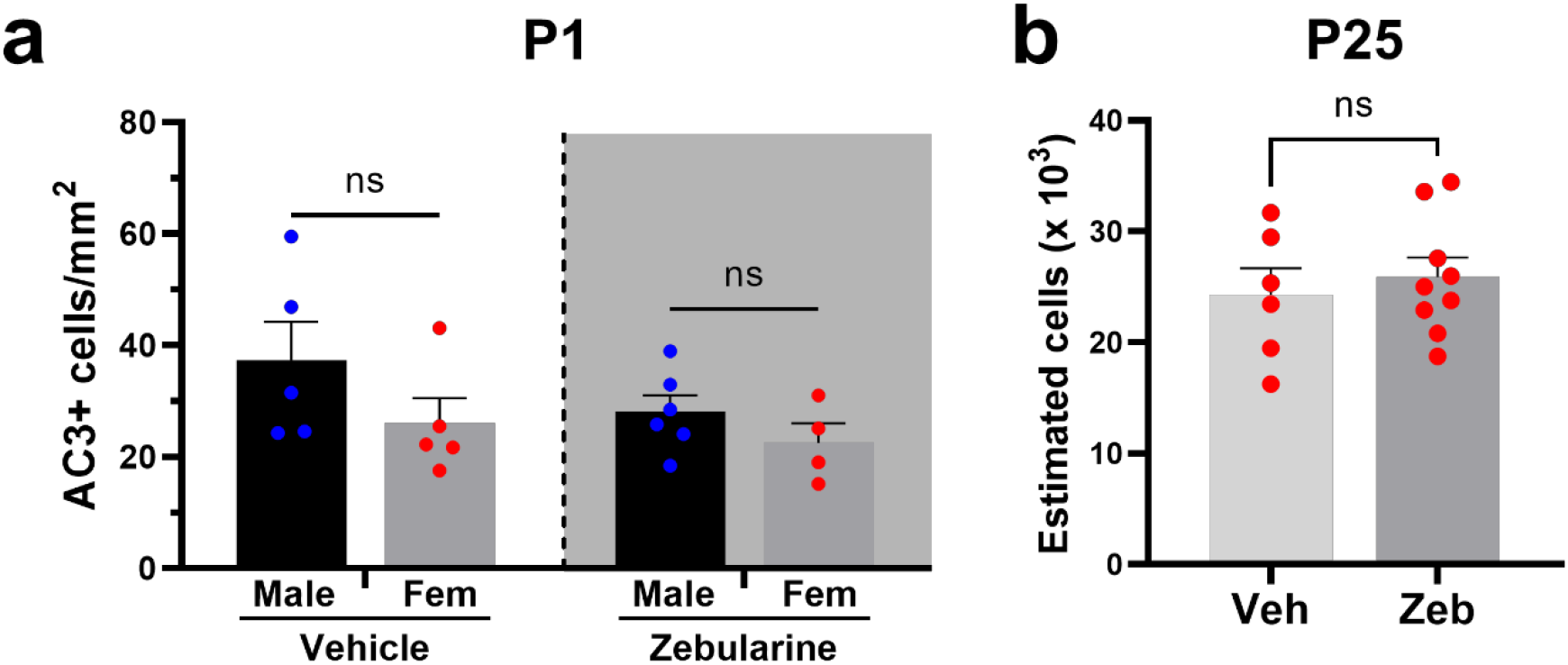
Zebularine treatment does not alter cell death on P1 or total cell counts in the ARC at weaning. (A) Number of cells in the ARC that were positive for activated caspase-3 (AC3) in males and females treated with vehicle or zebularine on P0 and P1. Animals were sacrificed 3h after the last injection. (B) Total neuron number in the ARC of P25 females that were treated with vehicle or zebularine neonatally. Mean + standard error of the mean and individual data points are depicted.

The results of this experiment were somewhat surprising because neonatal zebularine treatment eliminates the sex difference in ERα in the VMHvl by increasing expression specifically in males and testosterone-treated females, with no effect in females [14,16]. That is, a neonatal inhibition of DNA methylation leads to high, female-like ERα expression across groups in the VMHvl [16], but uniformly low, male-like ERα expression across groups in the neighboring ARC of the same animals. This suggested that DNA methylation regulates ERα expression very differently in these neighboring brain regions.

### Opposite effects of sex on DNA methylation of *Esr1* in the ARC and VMHvl

We used bisulfite conversion of DNA followed by pyrosequencing to compare effects of age, sex, and zebularine treatment on total methylation (5mC + 5hmC) of the ERα gene, *Esr1*, in the ARC and VMHvl. We focused on 16 CpG sites in two promoter regions shown to regulate expression of *Esr1* in the mouse brain: CpGs 1-11 in untranslated Exon A and CpGs 40-44 in untranslated Exon C [36,37].

The levels of total methylation of *Esr1* in the ARC and VMHvl observed at P1 were similar to those reported by Westberry et al. [36] in the neonatal mouse cortex using the same technique. In Exon A, repeated measures ANOVA indicated no overall sex difference in total methylation across the 11 CpG sites in either the ARC or VMHvl on P1 (Fig. 4a). However, sex differences in both brain regions emerged at P25, and in opposite directions: total methylation levels across the 11 CpG sites was significantly higher in females than males in the ARC (main effect of sex, F_1, 16_ = 8.63, *P* < 0.01), and higher in males than in females in the VMHvl (F_1, 16_ = 7.69, *P* < 0.015; Fig. 4a). Analyses of individual CpG sites showed that the pattern (F > M in the ARC, and M > F in the VMHvl) was consistent across most CpG sites (Fig. 4b). However, the difference reached significance only for CpG 10 in the ARC, and CpGs 6, 9, 10, and 11 in the VMHvl. The effects of sex were also specific to Exon A, as we found no sex differences in Exon C at P1 or at P25 in either the ARC or VMH (all *P*-values > 0.05; Suppl. Fig. 1).

**Fig. 4.**
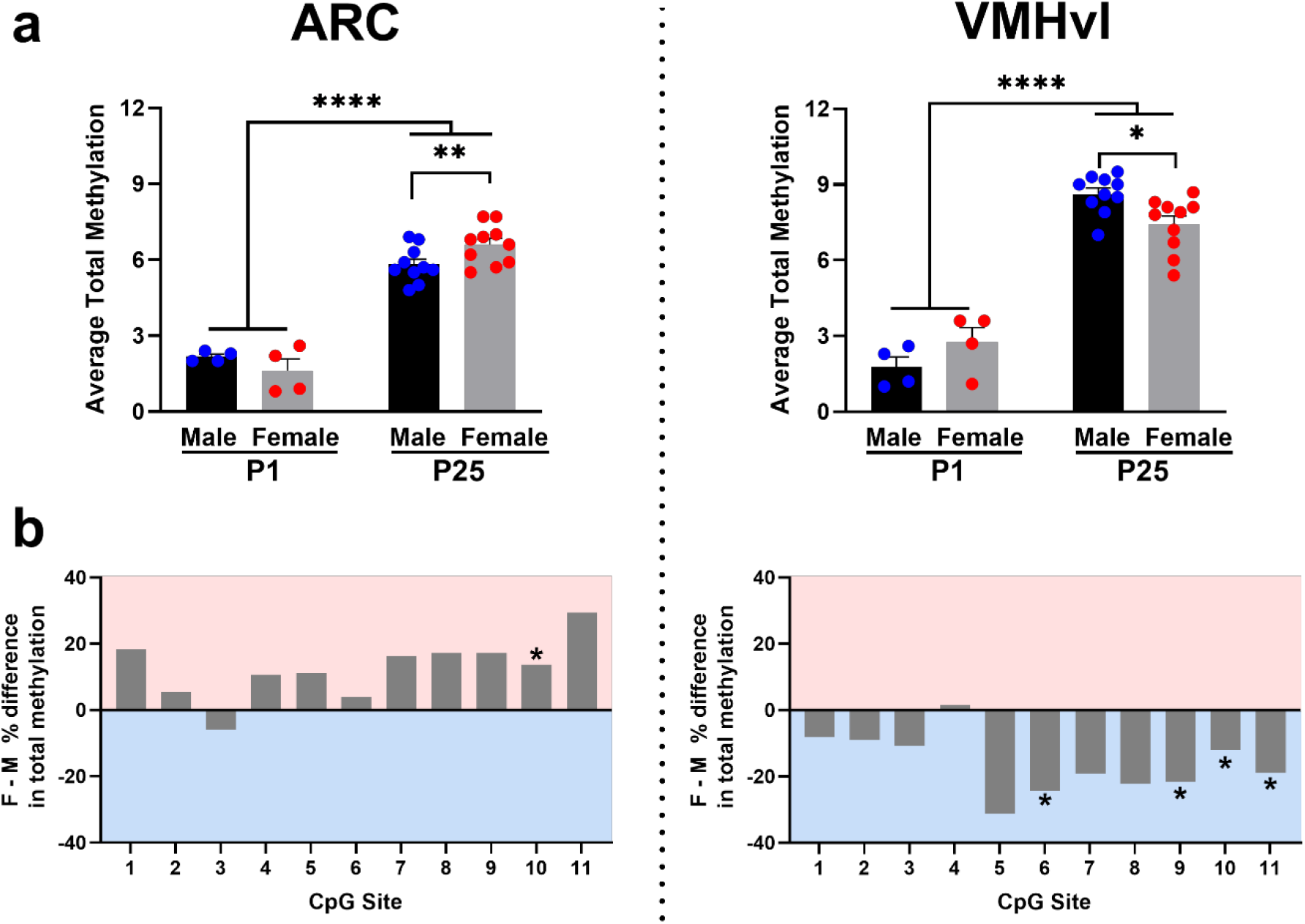
Sex differences in total DNA methylation of the *Esr1* Exon A promoter emerge at P25, and in opposite directions in the ARC and VMHvl. (A) Average total percent methylation across all 11 CpG sites in Exon A of the *Esr1* promoter in the ARC (left) and VMHvl (right) at P1 and P25 in male and female mice. Overall methylation increased markedly in both brain regions from P1 to P25. There were no significant sex differences at P1. At P25, total methylation was higher in females than in males in the ARC and higher in males than in females in the VMHvl. Mean + standard error of the mean and individual data points are depicted. (B) The sex difference in total methylation for individual CpG sites of Exon A at P25. Female minus male percent difference is plotted, e.g., [(Female % methylation – Male % methylation) / Male % methylation x 100), so that a difference of +20 means 20% more methylation in females at that site. In the ARC (left), most CpG sites had greater total methylation in females, although this was significant only for CpG 10 in posthoc tests (asterisk). In the VMHvl, most CpG sites had greater total methylation in males, and this was significant for CpG 6, 9, 10 and 11 in posthoc tests (asterisks).

The sex differences that emerged at P25 were superimposed upon large increases in average total methylation of CpG sites of the Exon A promoter region in both sexes and regions between P1 and P25 (main effect of age; F_1, 24_ > 200, *P* < 0.0001 in each region; Fig. 4a), consistent with our previous observation of an overall increase in global 5mC between birth and P25 in the mouse hypothalamus [30]. In addition, we found an interaction between age and sex in both regions (ARC: F_1, 24_ = 5.01, *P* = 0.035; VMH: F_1, 24_ = 7.59, *P* = 0.011), due to the fact that females gained more methylation with age than males in the ARC, and males gained more methylation with age than females in the VMHvl. There was a smaller increase in total methylation with age in Exon C when collapsing across sex (main effect of age; ARC: F_1, 24_ = 5.41, *P* = 0.029; VMHvl: F_1, 24_ = 6.52, *P* = 0.017; Suppl. Fig. 1). Paradoxically, we also found that neonatal zebularine treatment increased total methylation in Exon A at P25 in the ARC (F_1,16_ = 10.43, *P* < 0.01; Suppl. Fig. 2), with no sex-by-treatment interaction. Post hoc tests indicate that the effect was significant for CpG sites 6 and 9 (not shown). There was no effect of neonatal zebularine treatment on total methylation in the VMHvl.

### Down-regulation of *Tet* expression increases ERα in the ARC of males

The brain exhibits dynamic changes in DNA methylation during development [29,38,39], indicating an active methylation/demethylation cycle. TET enzymes control the turnover of DNA methyl marks, and mRNA expression of *Tet2* and *Tet3* is higher in neonatal males than in females in the POA and mediobasal hypothalamus [30]. This suggests that de-methylation may play a role in the development of sex differences. To test this, we administered siRNAs targeting *Tet2* and *Tet3* to newborn males, while control males and females received injections of non-targeting RNAs. *Tet2*/*Tet3* expression was significantly reduced in punches of the anterior and posterior hypothalamus 48 h after siRNA injection (main effect of siRNA, *Tet2:* F _1,10_ = 6.72, *P* = 0.027; *Tet3*: F_1,10_ = 5.21, *P* < 0.05; Suppl. Fig. 3). The reduction was fairly subtle (∼15-30%), but comparable to the magnitude of sex differences in *Tet* expression seen previously [30]. To determine whether this partial knock-down was functionally significant, ERα labeling was examined at P25.

In the ARC, we found a significant difference in ERα labeling across the three groups (F_2,20_ = 4.66, *P* < 0.025) and, as expected, females had greater ERα labeling than control males (*P* < 0.04). In males with *Tet2/Tet3* knock-down, ERα labeling was significantly increased relative to control males (*P* < 0.015), to a level very similar to that of females (*P* = 0.97; Fig. 5). In contrast, siRNAs against *Tet2 and Tet3* had no effect on ERα labeling in the VMHvl, although we did find the expected sex difference, with greater ERα labeling in females (F_2,21_ = 14.94, *P* < 0.0001).

**Fig. 5.**
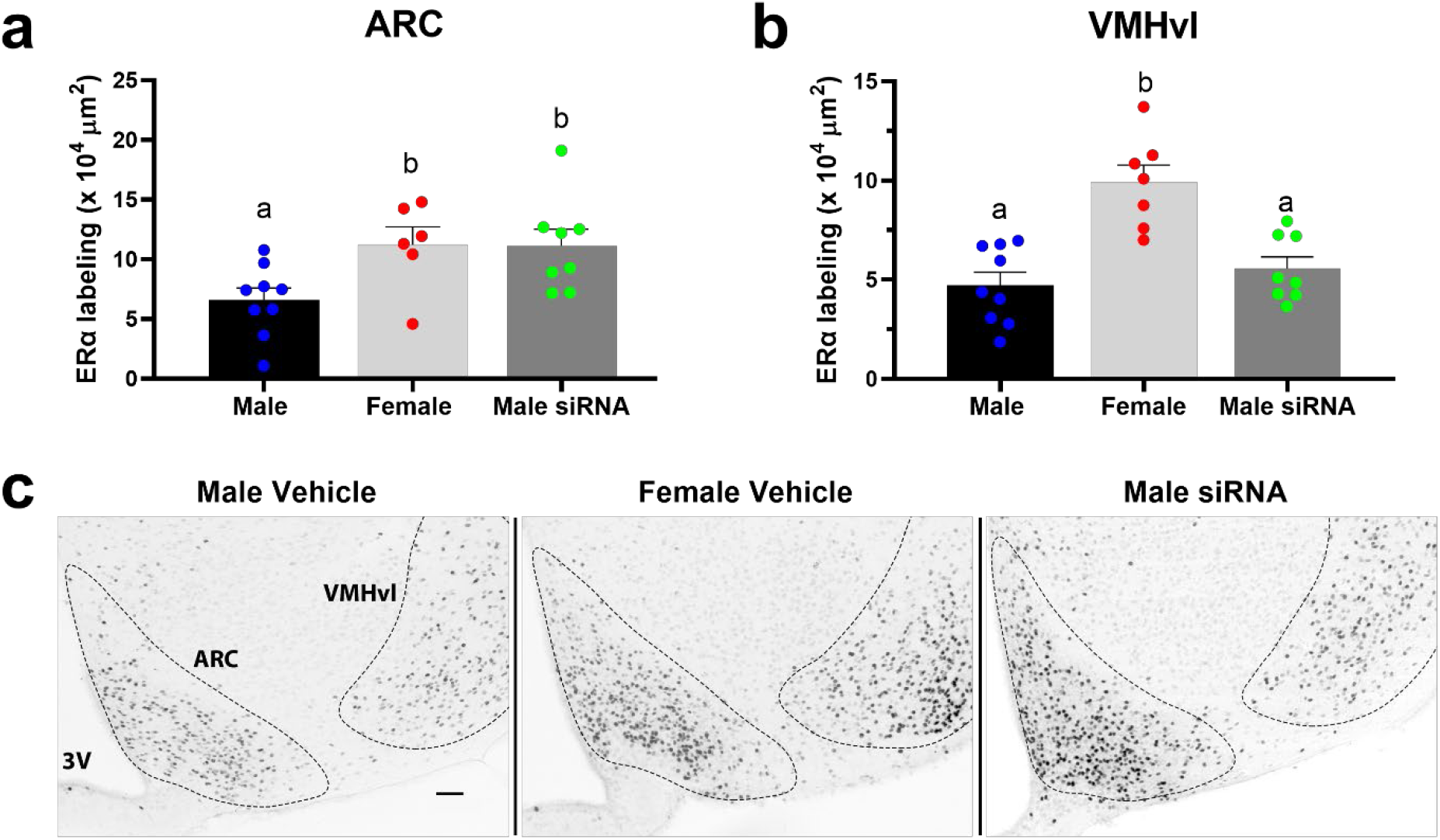
Neonatal downregulation of *Tets* abolishes sex differences in ERα labeling in the ARC, but not VMHvl. Mean ERα labeling at P25 in control males and females and in males with neonatal *Tet2* and *Tet3* knock-down in the (A) ARC and (B) VMHvl. Bars marked by different letters are significantly different (*P* < 0.05). (C) Photomicrographs showing ERα labeling in the ARC and VMHvl in the three groups. Scale bar = 50µm.

## Discussion

ERα neurons are highly enriched in the ARC and VMHvl, and play important roles in estradiol-mediated effects on energy homeostasis. A knockdown of *Esr1* in the ARC increases food intake and bone density in female mice [10,11], whereas blocking ERα signaling in the VMHvl reduces energy expenditure and thermogenesis [11,40,41]. In both regions, effects are sex-specific, with little or no effect of *Esr1* knockdown on energy homeostasis in males [9–11,42]. Females have more ERα neurons in both the ARC and VMHvl, and a similar female bias in ERα cell number has been reported in a variety of brain regions and vertebrate species [43–47]. In several cases, the sex difference requires differential exposure to testosterone during perinatal life [48–50], but the mechanism(s) by which testosterone programs sex differences in ERα expression, or other sex differences in neurochemistry for that matter, remains largely unknown.

We previously reported that sex differences in ERα in the POA and VMHvl of mice develop postnatally and are epigenetically regulated [14,30]. Both sexes express high levels of ERα at birth, and ERα cell number decreases in males over the next few weeks. The decrease can be prevented by treating neonatal males with an inhibitor of DNA methyltransferases, without an effect on cell death or total cell number [14]. We showed here that a sex difference favoring females also develops postnatally in the ARC and is abolished by inhibiting DNMTs during the first two days of life. However, in contrast to the POA and VMHvl, DNMT inhibition eliminated the sex difference in the ARC by decreasing (*i*.*e*., masculinizing) ERα expression in females.

We also found sex differences in total methylation of a promoter region of *Esr1* that emerged at weaning. Across 11 CpG sites of Exon A, males had a higher percent of total methylation than females in the VMHvl at P25. This is consistent with the hypothesis that 5mC marks contribute to the lower ERα labeling in the VMHvl of males. In the ARC, however, females had a higher percent of total methylation across Exon A, despite their higher levels of ERα. In addition, neonatal inhibition of DNMTs reduced ERα expression in the ARC of females, and neonatal knock-down of *Tet2/Tet3* expression in males, which is expected to increase 5mC, increased ERα cell number at weaning. Taken together, these observations suggest that 5mC marks across the CpG sites of Exon A are associated with *Esr1* gene activation in the ARC. Although contrary to the canonical association of DNA methylation with gene repression, there are a growing number of cases where DNA methylation of specific gene regions promotes transcription [20– 25].

One limitation of our findings is that pyrosequencing of sodium bisulfite-treated DNA does not distinguish 5mC marks from 5hmC. The concern in this case is lessened by the fact that 5hmC in neurons primarily accumulates in gene bodies and is depleted in promoter regions [29,35]. In addition, 5mC is 4-5 times more abundant than 5hmC in CpG dinucleotides [51]. Because we examined promoter regions and modifications in the CpG context, most of the total methylation we detected was presumably 5mC. Nonetheless, one would ideally like to sequence using techniques that distinguish 5mC from 5hmC [52], although the amount of starting material required is a serious impediment for this type of analysis.

The sex differences we detected in total methylation of *Esr1* were small (on the order of 2% in absolute terms and 20% relative differences between sexes across the CpG sites of Exon A), but similar in magnitude to what was previously reported for steroid receptors in the rat POA and mediobasal hypothalamus [39]. In our study and related previous studies [36,39,53], pyrosequencing of *Esr1* was examined from brain punches, which contain many cells types. ERα neurons comprise only a minority of all cells, even in regions such as the POA, VHM and ARC, where they are relatively abundant. Any “signal” (*i*.*e*., a sex difference in methylation of *Esr1*) must therefore be detected over quite a bit of noise. In addition, any measure of DNA methylation provides evidence that is correlative in nature. To test whether any given methyl mark *causes* a sex difference in ERα cell number would require specifically manipulating that mark, and while this is now theoretically possible [54,55], it may be some time before this can be achieved site-specifically, *in vivo*, in a newborn mouse brain.

In addition to the sex differences in total methylation that emerged at P25, we found that both sexes accumulate modified cytosines in Exon A, and to a lesser degree in Exon C, of *Esr1* from birth to weaning. This is consistent with previous findings of marked increases in 5mC and 5hmC throughout the genome in the mouse hypothalamus and neocortex from birth to adolescence [29,30]. The fact that females have persistently high levels of ERα in both the ARC and VMHvl despite the accumulation of methylation in a brain-relevant *Esr1* promoter, suggests that there are other regulatory mechanisms that allow for gene expression across development.

We expected zebularine, which inhibits DNMTs, to decrease 5mC marks and, therefore, total DNA methylation of *Esr1*, but did not observe that in either the VMHvl or ARC. In the VMHvl, there was no effect of neonatal zebularine treatment on total methylation of Exon A or Exon C, even though this treatment previously led to a lasting increase in ERα in males [14,16]. This suggests that zebularine may act indirectly (*e*.*g*. by inhibiting an inhibitor of *Esr1* in males), or that the *Esr1* promoter regions we examined do not capture the direct effects of zebularine. Although Exons A and C are important for developmental regulation of ERα in the brain, other promoters regulate ERα in other tissues [56], and methylation in non-promoter regions can also regulate gene expression [24,57,58]. Even more surprising, neonatal zebularine treatment increased the percent total methylation in Exon A of *Esr1* in the ARC at P25. If 5mC indeed activates *Esr1* in the ARC, the increased total methylation after zebularine would be consistent with the increased ERα seen in testosterone-treated females that received zebularine. However, the effect of zebularine was seen across all groups, which is harder to explain. We note that in *post hoc* analyses, the individual CpG sites affected by zebularine did not overlap with those that showed a significant sex difference in the ARC. In the current study, for example, CpG-10 was significantly different by sex, but not by zebularine treatment, in both the ARC and VMHvl. Differences in *Esr1* expression between peripheral tissues have been attributed to the methylation status of a single CpG site [59], and the effects of maternal behavior on glucocorticoid receptor gene expression in the brains of offspring have also been linked to a single CpG [60].

DNA demethylation and 5hmC marks are just beginning to be explored in the context of sexual differentiation. Given the unusually high abundance of 5hmC in gene bodies of neurons [61], it is likely to have an important role in shaping neural gene expression. We recently found more consistent sex differences in the expression of *Tets* than of *Dnmts* in the neonatal mouse brain [30], with higher *Tet2* and *Tet3* expression in males. The fact that suppressing *Tet2* and *Tet3* expression in males increased ERα in the ARC to female-like levels indicates that this sex difference is functionally meaningful. Thus demethylation, or possibly stable 5hmC marks, likely contribute to sexual differentiation of the brain.

Overall, our data suggest that neurons with the potential to express ERα are normally inhibited from doing so by the presence of DNA methyl marks in the VMH and the absence of such marks in the ARC. Since sex differences in both regions are due to perinatal gonadal steroids, and can be quite stable [62], hormones likely program ERα cell number through epigenetic mechanisms. In cancer cells, total methylation levels are reduced in response to estradiol [63], and one mechanism for this hormone-dependent hypomethylation is the upregulation of *Tet2* expression [64]. Neonatal estradiol also dampens the catalytic efficiency of DNMTs in the POA of the rat brain [12]. Thus, there may be parallel mechanisms by which sex steroid hormones influence DNA methylation, leading to stable changes in neurochemistry in the developing brain.

## Acknowledgements

We are appreciative of Taylor Hite for technical assistance and to members of the Forger Lab for feedback on the manuscript.

## Statement of Ethics

All procedures were performed in accordance with the National Institutes of Health animal welfare guidelines and were approved by the Georgia State University Institutional Animal Care and Use Committee.

## Conflict of Interest Statement

There is no conflict of interest regarding this work.

## Funding Sources

This work was supported by a National Science Foundation Graduate Research Fellowship awarded to LRC, and a National Institutes of Health Grant R01 MH068482 and Brains & Behavior Seed Grant from Georgia State University awarded to NGF.

## Author Contributions

LRC, CC, and NGF conceptualized the experiments. LRC, INKVC, and NGF wrote the manuscript. LRC, CC, INKVC, DM, EKR, and ACR performed data collection and/or analysis. All authors edited and approved the manuscript.

## Figure Legends

**Suppl. Fig. 1. There were no sex differences in average total methylation in Exon C of *Esr1*, but there was a subtle effect of age**. Average total methylation across 5 CpG sites in Exon C of the *Esr1* promoter in the ARC (left) and VMHvl (right) at P1 and P25 in male and female mice. There were no significant sex differences at P1 or P25 in the ARC (left) or VMHvl (right).

**Suppl. Fig. 2. Zebularine increases total methylation in Exon A of *Esr1* in the ARC**. Average total methylation across 11 CpG sites in Exon A of the *Esr1* promoter in the ARC was calculated for each animal. Zebularine-treated males and females had greater total methylation than vehicle-treated animals.

**Suppl. Fig. 3. Confirmation of *Tet2* and *Tet2* knock-down 48 hours after siRNA intracerebroventricular injection**. There was a modest, but significant reduction in *Tet2* (top) and *Tet3* (bottom) expression in tissue punches from the anterior and posterior hypothalamus (main effect of siRNA treatment, with no interaction).

## Notes

### Competing Interest Statement

The authors have declared no competing interest.

