## Supplementary figures and images for "DNA methylation and demethylation underlie the sex difference in estrogen receptor alpha in the arcuate nucleus"

### Supplemental Figure 1

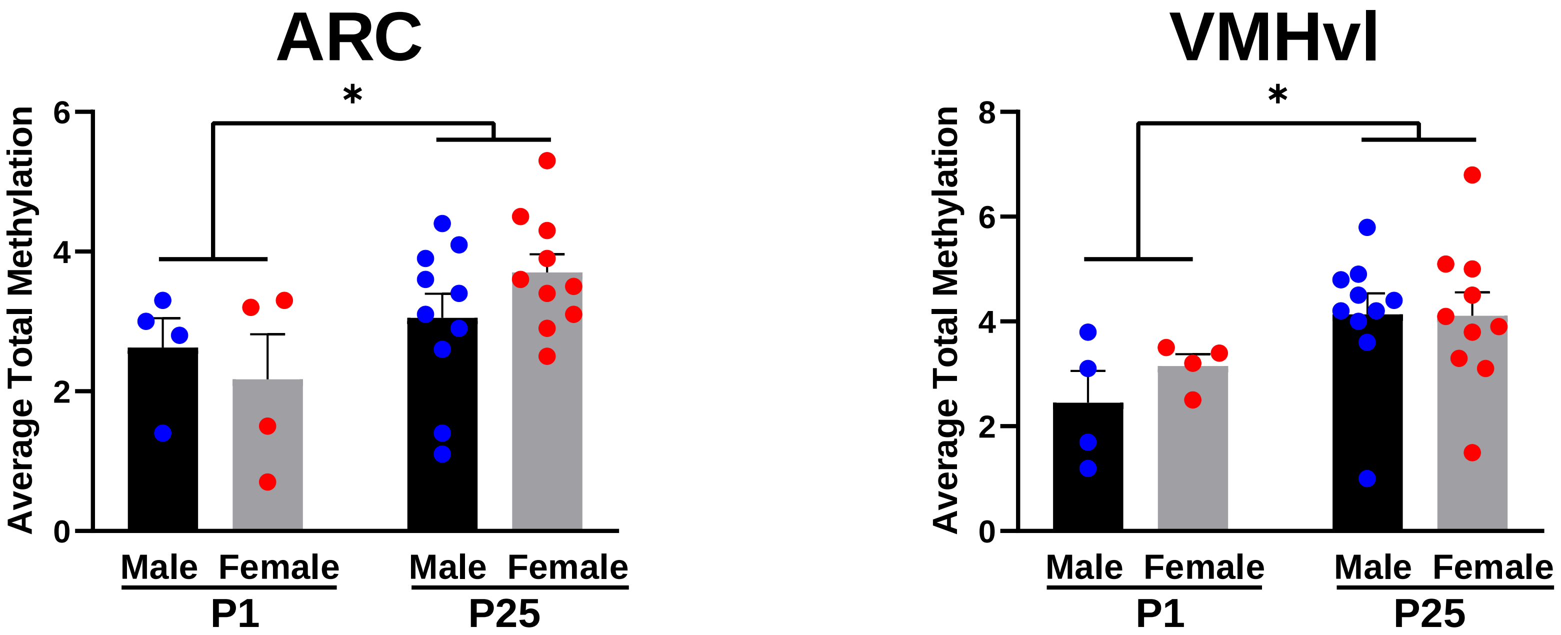

### Supplemental Figure 2

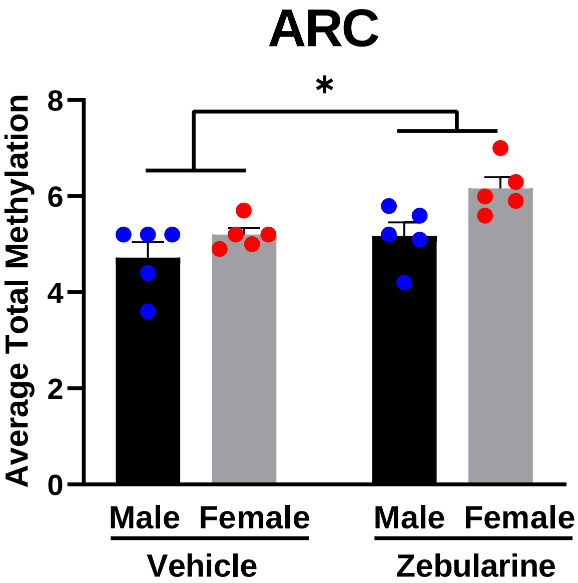

### Supplemental Figure 3

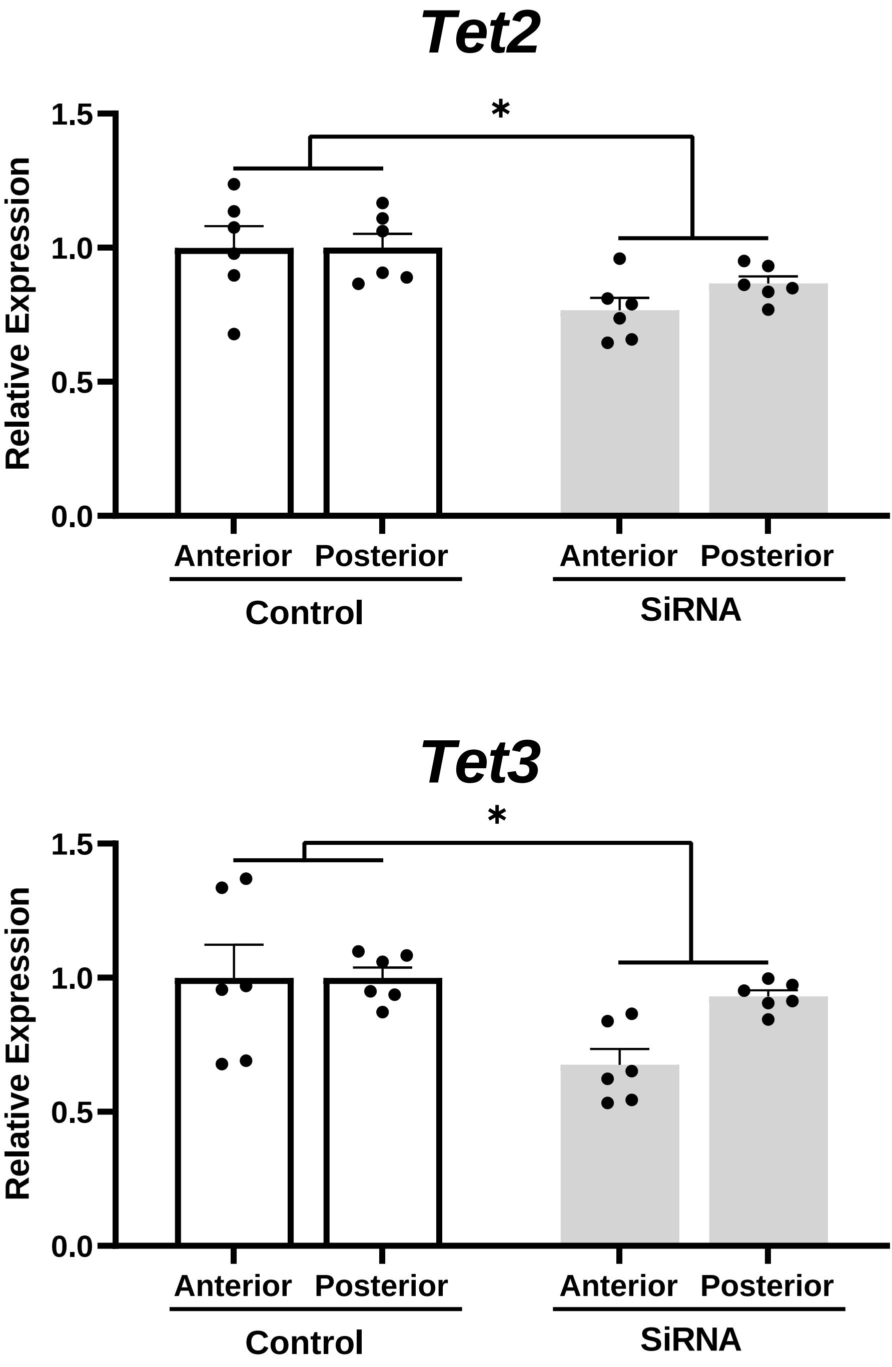
